# Sulfate limitation increases specific plasmid DNA yield and productivity in *E. coli* fed-batch processes

**DOI:** 10.1101/2023.02.09.527815

**Authors:** Mathias Gotsmy, Florian Strobl, Florian Weiß, Petra Gruber, Barbara Kraus, Juergen Mairhofer, Jürgen Zanghellini

## Abstract

Plasmid DNA (pDNA) is a key biotechnological product whose importance became apparent in the last years due to its role as a raw material in the messenger ribonucleic acid (mRNA) vaccine manufacturing process. In pharmaceutical production processes, cells need to grow in the defined medium in order to guarantee the highest standards of quality and repeatability. However, often these requirements result in low product titer, productivity, and yield.

In this study, we used constraint-based metabolic modeling to optimize the average volumetric productivity of pDNA production in a fed-batch process. We identified a set of 13 nutrients in the growth medium that are essential for cell growth but not for pDNA replication. When these nutrients are depleted in the medium, cell growth is stalled and pDNA production is increased, raising the specific and volumetric yield and productivity. To exploit this effect we designed a three-stage process (1. batch, 2. fed-batch with cell growth, 3. fed-batch without cell growth). The transition between stage 2 and 3 is induced by sulfate starvation. Its onset can be easily controlled via the initial concentration of sulfate in the medium.

We validated the decoupling behavior of sulfate and assessed pDNA quality attributes (supercoiled pDNA content) in *E. coli* with lab-scale bioreactor cultivations. The results showed an increase in supercoiled pDNA to biomass yield by 33 % and an increase of supercoiled pDNA volumetric productivity by 13 % upon limitation of sulfate.

In conclusion, even for routinely manufactured biotechnological products such as pDNA, simple changes in the growth medium can significantly improve the yield and quality.

**Highlights:**

- Genome-scale metabolic models predict growth decoupling strategies.
- Sulfate limitation decouples cell growth from pDNA production.
- Sulfate limitation increases the specific supercoiled pDNA yield by 33 % and the volumetric productivity by 13 %.
- We propose that sulfate limitation improves the biosynthesis of over 25 % of naturally secreted products in *E. coli*.

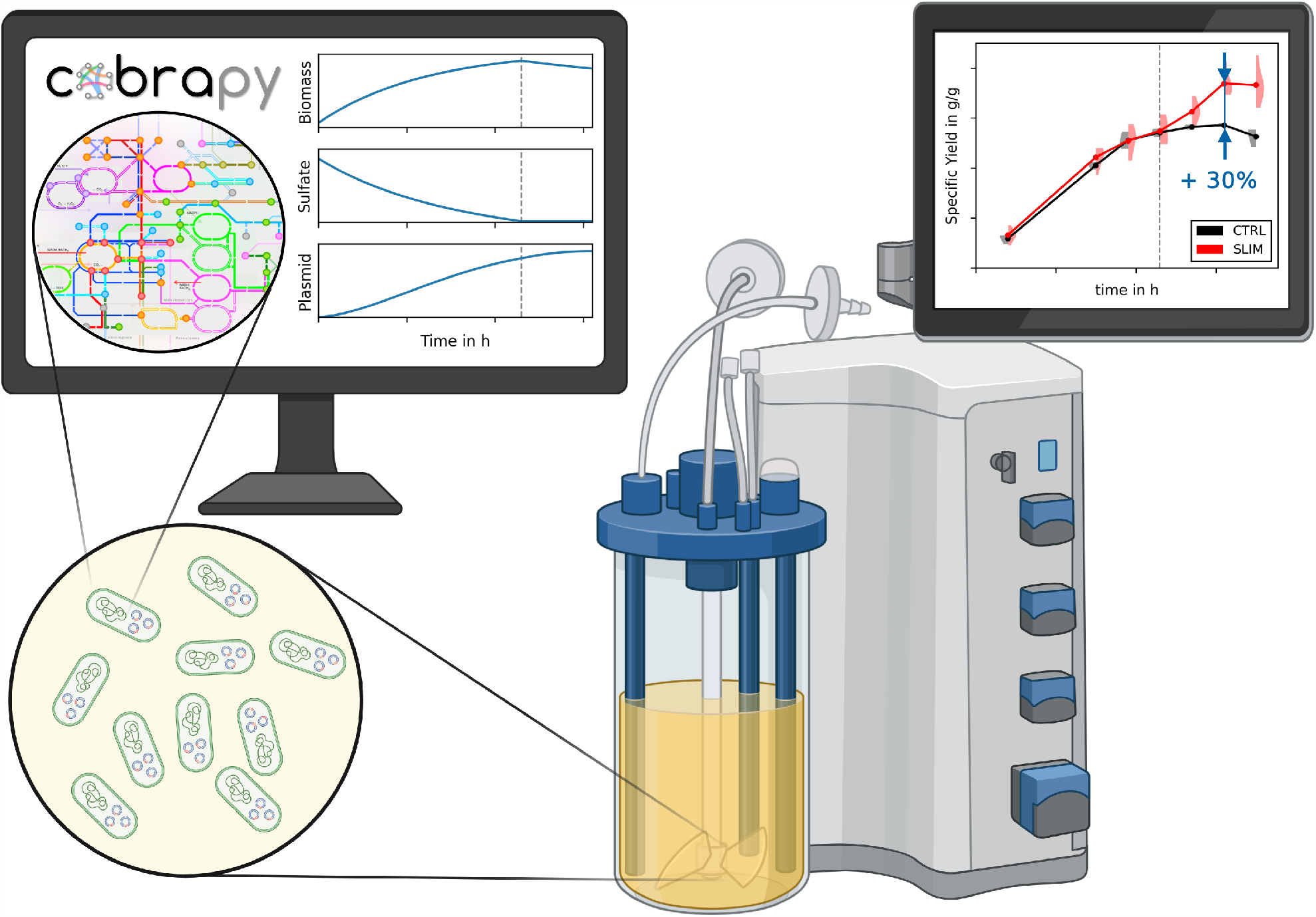

## 1. Introduction

Plasmid DNA (pDNA) is an important product of the pharmaceutical industry being primarily used as vectors for the transfection of mammalian cells. For example, pDNA can be directly injected in the form of a DNA vaccine [1]. Moreover, it is an important raw material for the production of mRNA vaccines, for example against SARS-CoV-2 [2, 3]. Additionally, pDNA can be used as a vector for gene therapy [4]. Regardless of the application, high amounts and high quality of pDNA are needed and the optimization of its production is of health-economic interest as pDNA is a relevant driver of manufacturing costs.

Apart from solely maximizing the yield of pDNA, three prerequisites are required for the design of a pDNA production process. Firstly, pDNA can be present in an open circular (oc), linear (l), or covalently closed circular (ccc), i.e., supercoiled form. Generally, the ccc form is considered more favorable for transfection of mammalian cells and, therefore, the fraction of supercoiling is of importance [5, 6]. Secondly, a loss of plasmid can be severely detrimental to the productivity during the fermentation process. Classically, this problem can be mitigated by the introduction of antibiotic resistance selection systems and the usage of antibiotic selection pressure during fermentation. However, these systems have several downsides. They, on the one hand, shift metabolic resources from the production of pDNA to the production of antibiotic resistance proteins [7]. On the other hand, special care has to be taken to remove residual antibiotics during pDNA purification and the absence thereof has to be validated. Therefore, several alternatives have been developed [8] although still not state-of-the-art yet for pDNA manufacturing. Thirdly, even though pDNA production in complex media generates higher yields, chemically defined media are preferred for the production of high-quality and safe pharmaceuticals [1].

Many strategies for the increase of pDNA production have been published with a large fraction using *E. coli* as production organism [9]. The methods range from the screening of favorable strains to metabolic engineering through knocking-in and -out of genes to antibiotic-free selection systems and other highly optimized production strains [9]. Most strategies for increasing pDNA production can be grouped into two conceptual approaches: (i) the reduction of cell growth; (ii) ensuring a constant supply of DNA precursor metabolites. The methods differ widely in the way one (or both) aims are achieved.

Early on researchers found that a low growth rate increases the specific productivity of pDNA [10]. To achieve this in batch fermentations Galindo et al. [11] designed a medium that releases glucose enzymatically and thus down-regulates glucose uptake and subsequently growth. Alternatively, Soto et al. [12] developed *E. coli* strain VH33 by knocking-out the main uptake pathway of glucose to achieve the same result. With this method, they could increase the production to 40 mg L^−1^ pDNA compared to 17 mg L^−1^ of the wild type strain [12]. Moreover, an optimization strategy for microaerobic environments was devised where *E. coli* strain W3110 improved pDNA production in presence of a recombinant expression of the *Vitreoscilla* hemoglobin protein [13]. In a subsequent study, this strain was tested in batch fermentations with different oxygen transfer rates and they concluded that as oxygen was depleted the growth rate decreased and the production of pDNA increased [14].

A strategy to ensure a constant supply of DNA precursor metabolites is, for example, the knocking-out of pyruvate kinase which forces metabolization of glucose over the pentose phosphate pathway [7, 15, 16]. Other methods utilize stoichiometric models to optimize the growth medium [17]. These authors concluded that the addition of the nucleosides adenosine, guanosine, cytidine, and thymidine as well as several amino acids can significantly improve pDNA production (60 mg L^−1^ in a batch fermentation). Martins et al. [18] optimized the growth medium for the high producer strain VH33 and concluded that the presence of aromatic amino acids (phenylalanine, tryptophan, tyrosine) is advantageous for redirecting molecules to the nucleotide synthesis pathways. Additionally, the effect of the amount and type of nitrogen source in the growth medium has been investigated on the production of pDNA to 213 mg L^−1^ [19]. Also, economical aspects of the medium design have been discussed [20].

Further potential for optimization is the pDNA itself. For example, reducing the size of the pDNA has been linked to higher volumetric yields [21]. However, a reduction might not always be possible, especially for therapeutic applications, where plasmid sizes are typically large (> 6 kb) [22]. Moreover, the pDNA yield of a process is highly dependent on the origin of replication. Currently, most plasmids carry a high copy number pUC origin of replication that allows up to 700 pDNA copies per cell [1]. Other approaches involved microaerobically induced [23] or heat-induced origins of replication that increase the plasmid copy number at higher temperatures than 37 °C [24, 25]. However, higher temperatures come with physiological trade-offs and, therefore, the amplitude and timing of heat induction are of importance [26].

Recently also other production organisms were proposed, e.g. *Lactococcus lactis*. Contrary to *E. coli, L. lactis* is generally regarded as safe (GRAS) and thus simplifies the downstream processing [27, 28].

Here, we design a three-stage bioprocess, where cellular growth and pDNA production are decoupled. We use constraint-based modeling to (i) identify medium components that induce the switching and (ii) determine the optimal time point for switching between the phases such that the average volumetric productivity is maximized.

## 2. Methods

### 2.1. Metabolic Modeling

All models and code used for the creation, simulation, and analysis of these models are available at https://github.com/Gotsmy/slim.

#### 2.1.1. Model Creation

For our analysis, we used *i*ML1515 [29], a genome-scale metabolic model of *Escherichia coli* strain K-12 substrain MG1655. All model modifications and simulations were performed in Python 3.10 using the CobraPy package [30]. To simulate plasmid production, an 8 base pair (bp) dummy pDNA metabolite with 50 % GC content was added to the model. Subsequently, we added a pDNA production reaction, corresponding to the dummy plasmid’s stoichiometry. pDNA polymerization cost was estimated to 1.36 mol adenosine triphosphate (ATP) per mol dNTP [7, 31]. Additionally, a pDNA sink reaction was introduced to make fluxes through the pDNA synthesis reaction feasible. An SBML version of the used model is available at https://github.com/Gotsmy/slim/tree/main/models.

Although 8 bp is an unrealistically small size for a real pDNA molecule, we emphasize that the actual length of the plasmid does not change the relative underlying stoichiometry. The reason we chose a small dummy plasmid is that it already has a molecular weight of approximately 4943.15 g mol^−1^. If we included a multiple kbp sized plasmid into the model, we would have risked that its large molecular weight lead to numerical instabilities during simulation [32]. However, all results are shown in gram pDNA, therefore, the exact length of the dummy plasmid does not change the values.

#### 2.1.2. Identification of Decoupling Compounds

Initially, we performed a parsimonious flux balance analysis (pFBA) [33] with biomass growth as objective. The maximal glucose uptake rate was set to 10 mmol g^−1^ h^−1^ and a non-growth associated maintenance requirement was set to 6.86 mmol ATP g^−1^ h^−1^ [29]. Exchange reactions with non-zero fluxes were used for the definition of the minimal medium. All exchange reactions that were not present in the minimal medium, except for H_2_O and H^+^, were turned off. To investigate the differences in uptake fluxes, an additional pFBA was performed with pDNA as the objective.

Next, we selected each of the minimal medium components and set the maximum exchange flux bound for this metabolite to 5, 25, 50, 75, and 100 % of the flux during biomass growth. For each value, a production envelope (pDNA synthesis as a function of growth) was calculated. Decoupling medium components were identified as metabolites which, as their uptake flux decreased, the maximum pDNA synthesis potential increased at a maximum biomass growth.

#### 2.1.3. Process Simulation

We used dynamic flux balance analysis (dFBA) [34] to simulate the time evolution of the fed-batch processes. Only the feed phase was simulated (stage 2 and 3 in Figure 1) as the batch remained unchanged. At every integration step, a lexicographic flux balance analysis (FBA), where all fluxes of interest were consecutively optimized, was performed [35]. The list of objectives is given in Table 1. The non-growth associated maintenance was kept at a constant value as before. We used SciPy’s `solve_ivp` function for numerical integration [36].

**Table 1.**
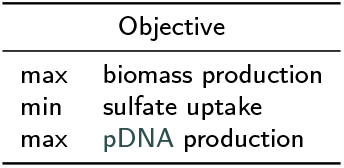
Lexicographic objectives for dFBA. The order of optimization was top to bottom.

**Figure 1:**
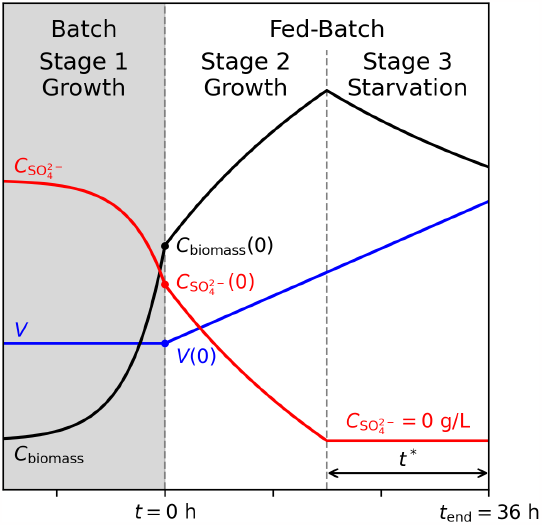
Schematic of a three-stage growth-decoupled fed-batch pDNA production process. (1. batch, 2. fed-batch with cell growth, 3. fed-batch without cell growth). Size of the variables and length of the stages are not to scale. As the batch process was kept unchanged, it was not included in the fed-batch optimization simulations. Instead, the simulation starts at the beginning of the feed (*t* = 0 h) with realistic values for the process variables at batch end (Table S1).

Sulfate concentration, 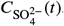, in the medium was tracked as its depletion coincides with the metabolic switch from biomass growth to production. Due to a lack of knowledge, no uptake kinetics of sulfate were simulated. Practically, this meant that the sulfate uptake was calculated from the biomass stoichiometry and growth rate. The exchange reaction bounds were left unconstrained when 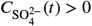 and were blocked when 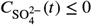.

High copy number origin of replication plasmids typically replicate during the growth phase as they high-jack genomic DNA synthesis pathways. Therefore, we set a lower bound, 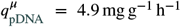, for the pDNA synthesis reaction. Moreover, it is unrealistic to assume that all available glucose is channeled towards pDNA production during the 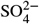 starvation. Therefore, we set an upper bound to the synthesis reaction. Since its actual value was unknown, we tested several levels ranging from 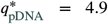 to 24.7 mg g^−1^ h^−1^. Throughout this manuscript, that ratio of upper to lower bound of the pDNA synthesis reaction is referred to as

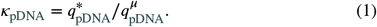

We simulated 41 equidistant levels of *κ*_pDNA_ ∈ [1, 5]. Because of the implementation of the dFBA, the lower and upper bounds of the synthesis reaction can be interpreted as the pDNA synthesis fluxes during the growth and 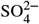 starvation phase, respectively. We assumed negligible changes in overall biomass composition due to pDNA synthesis, which solely derives from external carbon sources.

To compare pDNA production processes, we calculated two performance indicators: the specific yield

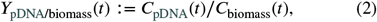

and the average volumetric productivity

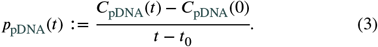

Here, *t*_0_ = 0 h indicates the start of feeding (see Figure 1). With our parameters, see Table S1, (3) simplifies to

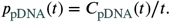

A schematic of a three-stage growth-decoupled fed-batch process is shown in Figure 1. Initial conditions (at the start of the feed phase) resembled realistic values from the end of a batch process in a small bioreactor (Table S1). Strategies with a linear (i.e. constant) and an exponential feeding rate,

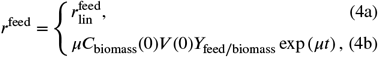

respectively, were tested. The specific glucose uptake rate was set to

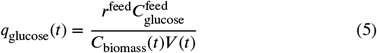

to ensure *C*_glucose_(*t*) = 0 throughout the (non-starved) fed-batch phase (stage 2, Figure 1). Here, 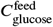 denotes the glucose concentration in the feed medium.

The dFBA simulation terminated once the maximal volume of 1 L was reached (i.e. after 35.2 and 36.0 h in exponential and linear fed-batch, respectively). The initial sulfate concentration in the medium at the start of feed was varied in 301 equidistant steps to search for a productivity optimum. We assumed that sulfate was present in the medium at the start of the feed, while the feed medium was sulfate-free.

#### 2.1.4. Identifying Alternative Targets

We screened 307 products with existing exchange reactions in *i*ML1515 [29] by performing lexicographic FBA (analogous to Table 1) with and without sulfate in the growth medium. Bioproducts for which the calculated synthesis rate improved during sulfatelimitation were identified as potential products benefiting from a sulfate limited process design.

### 2.2. Validation Experiments

#### 2.2.1. Upstream Process

All experiments were conducted with proline auxotroph *E. coli* K-12 strain JM108 [37], which previously had been used for pDNA production [8]. The cells were transformed with a plasmid of 12.0 kbp length and 53 % GC content containing a pUC origin of replication and a kanamycin resistance gene. However, no kanamycin was added throughout the production process as U.S. Food and Drug Administration (FDA) and European Medicines Agency (EMA) recommend to avoid the use of antibiotics [1].

For fed-batch fermentations, *E. coli* JM108 were grown in a 1.8 L (1.0 L net volume, 0.5 L batch volume) computer-controlled bioreactor (DASGIP parallel bioreactor system, Eppendorf AG, Germany). The bioreactor was equipped with a pH probe and an optical dissolved oxygen probe (Hamilton Bonaduz AG, Switzerland). The pH was maintained at 7.0 ± 0.1 by addition of 12.5 % ammonia solution; the temperature was maintained at 37.0 ± 0.5 °C. The dissolved oxygen (O_2_) level was stabilized above 30 % saturation by controlling the stirrer speed, aeration rate, and gassing composition. Foaming was suppressed by the addition of 2 mL 1:10 diluted Struktol J673A antifoam suspension (Schill+Seilacher, Germany) to the batch medium and by the automatic addition of 1:10 diluted Struktol J673A controlled by a conductivity-operated level sensor. For the inoculation of the bioreactor, a seed culture was used (25 mL batch medium inoculated with 250 μL master cell bank in 250 mL baffled glass flasks at 37 °C with shaking at 180 rpm). The seed culture was incubated until a final OD_600_ of 2-4 was reached and a defined volume was transferred aseptically to the bioreactor to result in an initial OD_600_ of 0.015.

The fermentation process was designed for a final amount of 50 g cell dry mass (CDM) of which 1.51 g was obtained in a batch volume of 500 mL and 48.5 g during the feed phase via the addition of another 500 mL of feed medium. The amount of glucose for the specific medium was calculated based on a yield coefficient (*Y*_biomass/glucose_) of 0.303 g g^−1^ and added as C_6_H_12_O_6_ · H_2_O. For media preparation, all chemicals were purchased from Carl Roth GmbH (Germany) unless otherwise stated. Two different media compositions were compared regarding specific pDNA productivity: one with a limited sulfur source (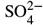 limitation) and one without a limited sulfur source (control). Feeding was initiated when the culture in the batch medium entered the stationary phase. A fed-batch regime with a linear substrate feed (0.26 g min^−1^ respectively 13.91 mL h^−1^) was used for 35 h (approximately five generations). During 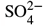 limitation fermentations, the simulations predicted that the provided sulfur was completely consumed at 23 h after feed start. The batch and fed-batch medium components are given in Table S2. The cultivations of 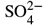 limitation and control were conducted in six and three replicates, respectively.

#### 2.2.2. Analysis

For off-line analysis (OD_600_, CDM, pDNA product), the bioreactor was sampled during the fed-batch phase. The OD_600_ was measured using an Ultrospec 500 pro Spectrophotometer (Amersham Biosciences, UK), diluting the samples with phosphate-buffered saline to achieve the linear range of measurement. For the determination of CDM, 1 mL of cell suspension was transferred to pre-weighed 2.0 mL reaction tubes and centrifuged for 10 min at 16.100 rcf and 4 °C with an Eppendorf 5415 R centrifuge. The supernatant was transferred to another reaction tube and stored at −20 °C for further analysis. As a washing step, the cell pellet was resuspended in 1.8 mL RO-H_2_O, centrifuged, and the supernatant discarded. Afterwards, the pellet was resuspended in 1.8 mL RO-H_2_O and finally dried at 105 °C for 24 h. The reaction tubes with the dried biomass were cooled to room temperature in a desiccator before re-weighing.

For pDNA product analysis, the sampling volume of the cell suspension, corresponding to 20 mg CDM, was estimated via direct measurement of the OD_600_. The calculated amount was transferred to 2.0 mL reaction tubes and centrifuged at 16.100 rcf and 4 °C for 10 min. The supernatant was discarded, and the cell pellets were stored at −20 °C.

The content of pDNA in ccc-conformation was determined using AIEX-HPLC (CIMac™ pDNA-0.3 Analytical Column, 1.4 μL; BIA Separations d.o.o., Slovenia). The column separated open circular, linear, and supercoiled pDNA fractions into distinct peaks. Quantification was achieved using a calibration curve based on peak areas obtained from purified pDNA samples. For HPLC analysis cell disintegration was performed by an alkaline lysis method [38]. The obtained lysate was directly analyzed by HPLC (Agilent 1100 with a quaternary pump and diode-array detector (DAD)). Values derived from three biological replicates have a coefficient of variation lower than 10 %.

The average volumetric productivity (*p*_pDNA_(*t*)) and the average specific yield (*Y*_pDNA/biomass_(*t*)) were calculated as given in Equations (3) and (2), respectively. The time-dependent pDNA synthesis rate (*q*_pDNA_(*t*)) was estimated via the `finitediff` Python package [39, 40].

## 3. Results

### 3.1. Key Objective

We aim to design an efficient three-stage fed-batch process (Figure 1) for pDNA production in *E. coli*, where cellular growth and pDNA production are separated.

In the following, we will

i. use constraint-based modeling to identify medium components that enable switching from growth to production phase;
ii. determine optimal switching time points to maximize the average productivity in a fed-batch fermentation; and
iii. experimentally validate the computed strategies in a linear fed-batch process.

### 3.2. Identification of Decoupling Compounds

First, we used pFBA to compute a minimal set of uptake rates in the genome-scale metabolic model *i*ML1515 [29] supporting maximal aerobic growth of *E. coli* with glucose as carbon source. Similarly, we computed uptake rates for maximal pDNA production using the same constraints (Figure 2). All calculated uptake rates are inflexible in the optima, except for Fe^2+^ and O_2_. Their uptake can be further increased by conversion to and excretion of Fe^3+^ and H_2_O. In contrast to biomass synthesis, pDNA production requires only glucose, O_2_, 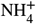, and 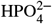, but no further nutrients. Therefore, we conclude that these remaining nutrients could potentially be used as decoupling agents separating pDNA synthesis from growth.

**Figure 2:**
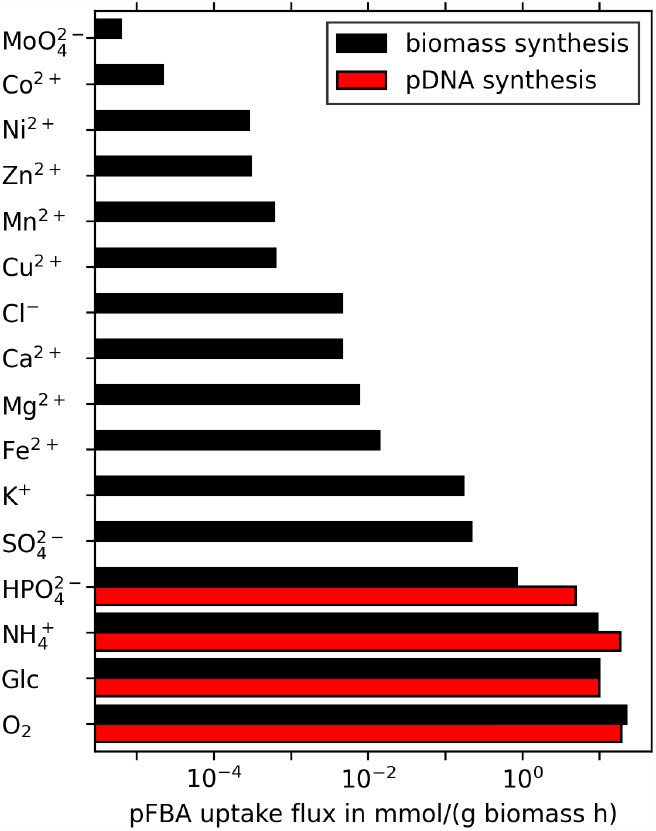
**pFBA uptake flux rates** for all minimal medium components. Black bars represent fluxes for optimization of biomass synthesis, red bars represent fluxes of pDNA synthesis.

Next, for each decoupling nutrient, we restricted its uptake between zero and 100 % of its rate at maximum growth and computed the corresponding pDNA production envelopes as a function of growth (Figure 3). All twelve decoupling nutrients result in identical sets of production envelopes, which mirrors the fact that each decoupling nutrient is essential for growth but not required for pDNA production. Note that the one-to-one trade-off between biomass production and pDNA synthesis, i.e., the upper limit of the production envelope, is a straight line between the points (0|100) and (100|0).

**Figure 3:**
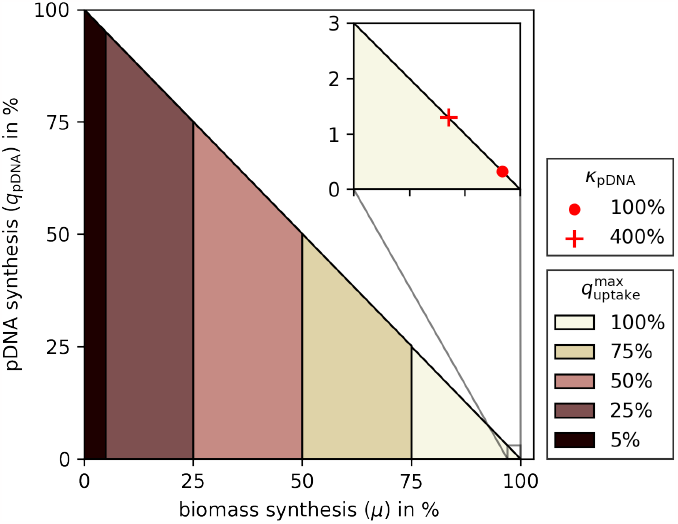
Normalized pDNA production envelopes for different maximal uptake rates of (a single) decoupling nutrient. Ca^2+^, Cl^−^, Co^2+^, Cu^2+^, Fe^2+^, K^+^, Mg^2+^, Mn^2+^, 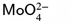, Ni^2+^, 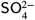, and Zn^2+^ all result in identical sets of production envelopes. Note that all production envelopes include the line segment from (0|0) to (0|100). The inset shows the extreme points of a production envelope with a realistic pDNA production rate during biomass growth (red circle) and a potential 4-fold pDNA production rate increase (red cross).

Realistically, pDNA production rates are significantly less than the theoretical value of 100 % in Figure 3. Therefore, in the inset of the same figure, the red markers illustrate the extreme points of more reasonable production envelopes. For example, the red circle (i.e., *κ*_pDNA_ = 100 %) illustrates average rates of pDNA production during cell growth. Even when the decoupling would lead to a boost by a factor of *κ*_pDNA_ = 4 (red cross), the resulting pDNA production flux would be only 1.3 % of the theoretical maximum.

In the following, we focus on the impact of the six bulk non-metal elements (sulfur, phosphorus, oxygen, nitrogen, carbon, and hydrogen) that typically make up 97 % (g g^−1^) of the elemental biomass composition [41]. Moreover, except for potassium (and sulfate), all other predicted decoupling nutrients (iron, magnesium, calcium, chlorine, copper, manganese, zinc, nickel, cobalt, and molybdenum) are taken up at minute rates (< 15 μmol g^−1^ h^−1^, Figure 2). Thus, exactly dosing their concentrations for limitation may be challenging in a bio-process. This leaves sulfate as the only predicted decoupling nutrient in a glucose-minimal medium.

### 3.3. Optimal Sulfate Limited Processes

Decoupling production from growth during a bio-process raises the question of timing: when to best switch from growth to production phase to maximize performance.

In the following, we used dFBA [34] to track the time-dependent concentrations *C*_*i*_(*t*) of biomass, pDNA, glucose, and sulfate and determine the optimal initial sulfate concentration, 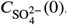, that maximize the average volumetric productivity

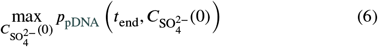

in a fed-batch process. Here *t*_end_ denotes the end of the bio-process, which terminates when the maximal volumetric capacity of the reactor is reached. Our simulations assume that pDNA production occurs (i) at a constant rate 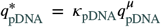 during the sulfate starvation phase; (ii) at 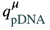 during the growth phase.

We simulated sulfate limited fed-batch processes with a linear feed as sketched in Figure 1 using the values listed in Table S1. In all simulations the feed rate was constant. Thus, the process length is always 36 h. Subsequently, we analyzed the impact of the length of the pDNA production phase (induced by sulfate starvation during the feed phase, i.e. stage 3 in Figure 1) on the average volumetric productivity. An analogous analysis for a sulfate limited batch fermentation can be found in the Supplementary Notes A.3.1.

We observed distinct maxima in the average volumetric productivity of a linear fed-batch when *κ*_pDNA_ > 1 (red dotted line, Figure 4A). Maxima occur at much longer starvation times compared to batch processes (*t*^*^ = 19 h versus 2.5 h at *κ*_pDNA_ = 4, Figure S1). Even compared to an equivalent exponential fed-batch, the optimal starvation is longer in a linear than in an exponential fed-batch process (Figure S2). For instance, at *κ*_pDNA_ = 1.5 a linear fed-batch process achieves an optimal *p*_pDNA_ = 0.18 g L^−1^ h^−1^ at *t*^*^ = 12 h, while an equivalent exponential fed-batch process reaches its optimum *p*_pDNA_ = 0.096 g L^−1^ h^−1^ at *t*^*^ = 3.3 h. Within our modelling assumptions, a linear fed-batch, even without starvation, outperforms an equivalent exponential fed-batch with (optimal) sulfate starvation as long as *κ*_pDNA_ ≤ 3.5.

**Figure 4:**
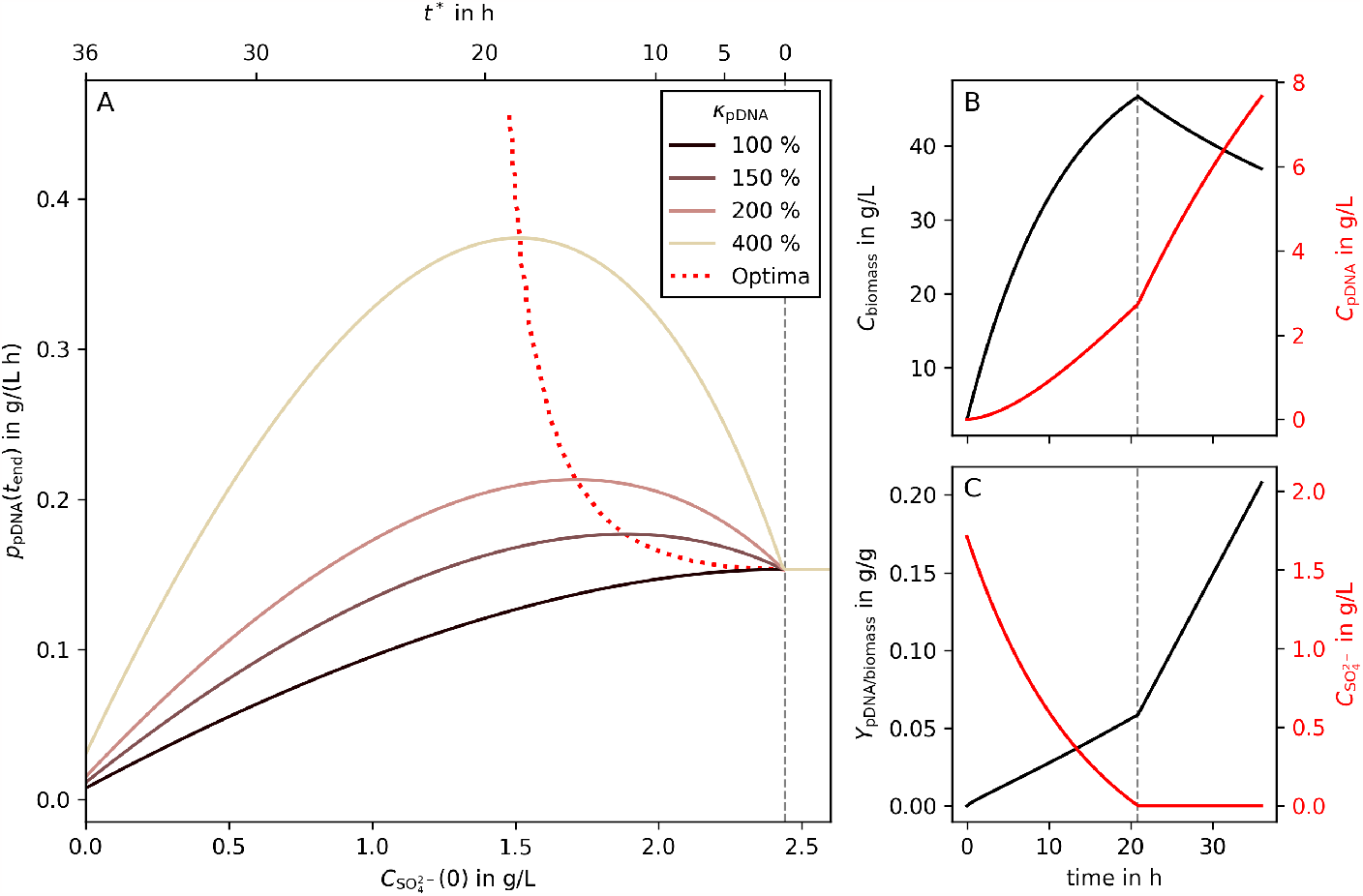
Predicted optimal timing in a 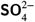 limited linear fed-batch process. Panel **A** shows average volumetric productivities of pDNA production (g L^−1^ h^−1^ 2−) as a function of the initial 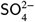 concentration in a linear fed-batch process. The full line represents different levels of increased pDNA production during starvation as percentages of pDNA production rate during biomass growth (*κ*_pDNA_). The dotted line indicates the location of the optima for *κ*_pDNA_ between 100 and 500 %. The second X-axis of panel **A** (top) illustrates the length of the sulfate starved process phase (*t*^*^). For all modeled linear fed-batch processes, right of the gray dashed line, no SO^2−^ limitation occurred. Panels **B** and **C** show the process curves of metabolites of interest in the optimal process for *κ*_pDNA_ = 200 %. The gray dashed lines indicate the switching time point between the growth and production phases.

Typically, growth-decoupled processes suffer from a substantially decreasing (glucose) uptake rate during the production phase [42, 43]. Thus, our assumption of keeping an elevated 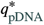 constant over several hours may be unrealistic. Therefore, we investigated how the productivity optima change when the maximal feasible starvation length 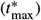 is bounded. Yet, even in such cases, sulfate limited fed-batches perform better than standard processes without starvation (Figure S3).

### 3.4. Sulfate Limitation Experiments

The preceding analysis suggested that a three-stage fed-batch process with sulfate starvation will deliver superior pDNA production performances compared to a conventional, non-starved fed-batch process. To confirm this, we set up a linear fed-batch process with *E. coli* JM108 as host (see Section 2.2 for details). Based on small molecule production rates during sulfate starvation [43, 44], we assumed a *κ*_pDNA_ = 2 and consequently predicted *t*^*^ = 13 h. Thus, we computed the initial 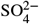 concentration to be 3.8 g L^−1^ such that sulfate starvation occurs after 23 h in a 36 h bio-process.

Figure 5 highlights the feasibility of sulfate starvation (indicated as SLIM) to boost pDNA production in a (linear) fed-batch. Panel A illustrates the growth arrest due to sulfate starvation (compare the diverging lines to the right of the dashed line). Due to dilution, the biomass concentration (red) decreases if cells no longer grow. Yet, pDNA concentration keeps rising – even faster than in the unstarved control (compare red SLIM with black CTRL in panel B). Consequently, the specific pDNA yield rises too (panel C) reaching a maximum of 0.074 g g^−1^ after 31 h, which corresponds to an improvement of 29 % compared to control (*p* = 0.0005, Table S3).

**Figure 5:**
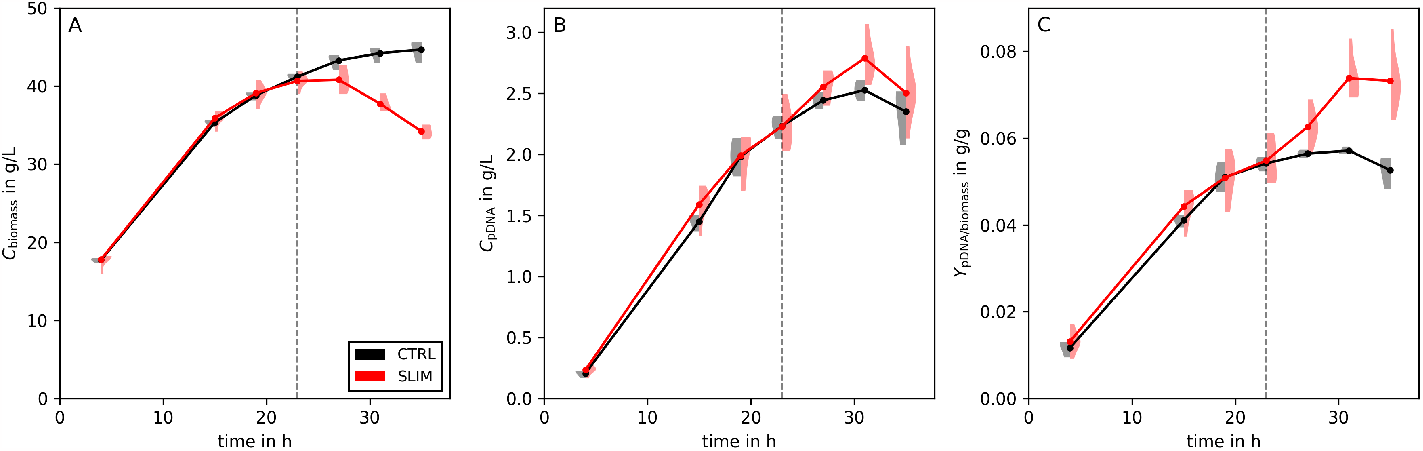
Experimental results of sulfate limitation. Panel **A** illustrates the biomass concentration, panel **B** the concentration of produced pDNA, and panel **C** the pDNA to biomass yield. The violins are calculated from triplicates of the control (i.e., no 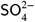 limitation, CTRL, black) and six replicates of sulfate limited processes (SLIM, red) are shown. The full lines and points are calculated from the mean of the replicates. The gray dashed line represents the estimated time of the switch from biomass growth to pDNA production (projected at 23 h).

Moreover, we compared the fraction of supercoiled pDNA between the 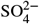 limited and control processes (Figure 6). Up to 27 h after induction, there were no noticeable differences. After this point, the 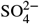 limited process maintained a higher fraction of supercoiled pDNA (+3%, *p* = 0.0019, Table S3), resulting in a 33 % (*p* = 0.0001) increase in the specific yield of supercoiled pDNA.

**Figure 6:**
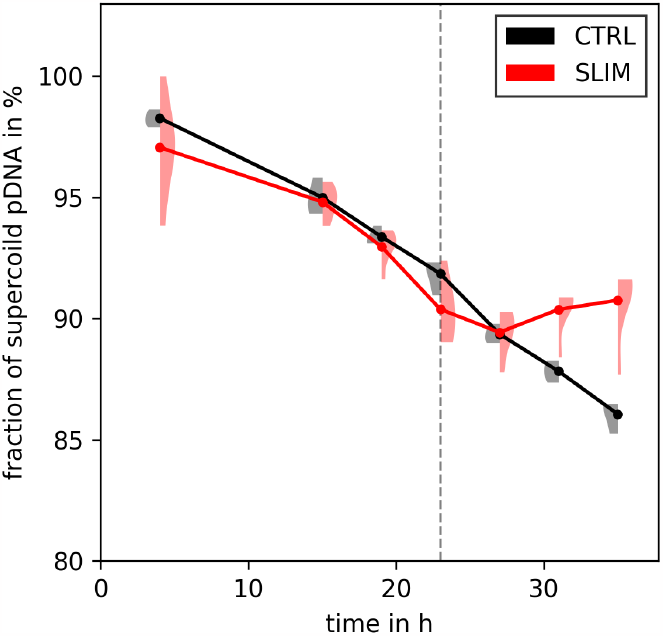
**Fraction of supercoiled (ccc) pDNA over time** for 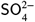 limited (SLIM, red) and control (CTRL, black) process. Experimental replicates are shown as violins, full lines and markers represent their means.

Beyond 31 h pDNA concentration and specific yield decrease in both the 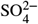 limited and the control process.

Next to the specific yield, also the average volumetric productivity increases by 10 % at 31 h (*p* = 0.0243, Figure 7A and Table S3). If only the pharmacologically relevant supercoild fraction of pDNA is considered, the productivity increases by 13 % (*p* = 0.0052, Table S3).

**Figure 7:**
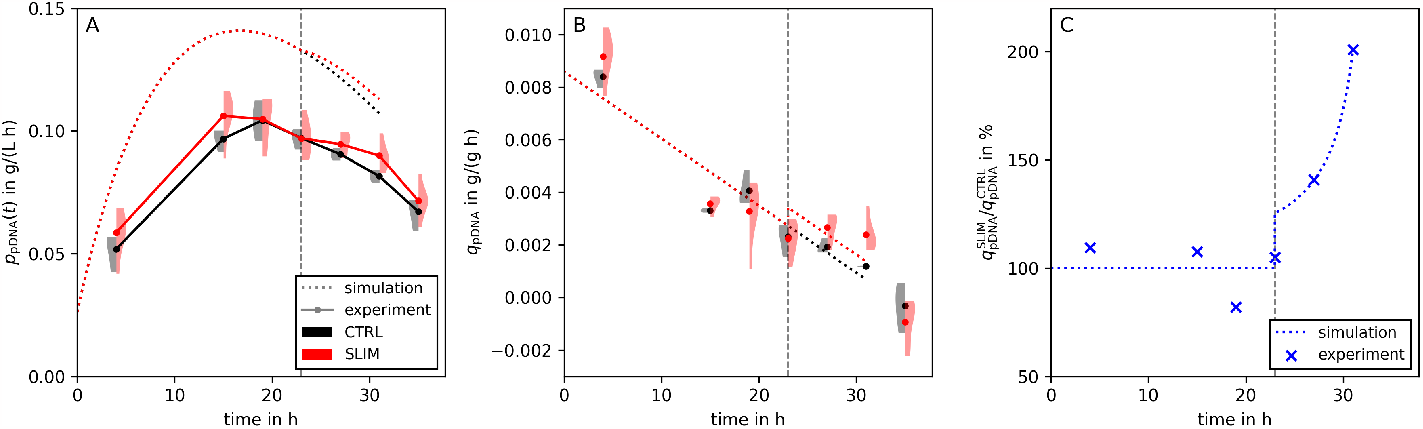
Calculated experimental rates and comparison to the simulation. Panel **A** illustrates the average volumetric productivity and panel **B** the pDNA production fluxes of control and 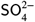 limited process (black CTRL and red SLIM, respectively). Panel **C** shows the ratio of 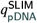 and 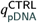 (blue). Experimental replicates are shown as markers, dotted lines are calculated from simulations. To adjust the simulations to the rates obtained in the experiments, a linearly decreasing 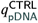 was fitted to experimental control data (black dotted line, panel **B**). Moreover, we fitted a parallel 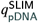 (red dotted line, panel **B**) to conform to the experimental flux ratio (panel **C**). The vertical gray dashed line represents the estimated time of switching from biomass growth to pDNA production (projected at 23 h).

To further investigate the experimental results, we computed the specific productivities in the sulfate limited and control fermentations (Figure 7B). In both bio-processes *q*_pDNA_ decreases with time which is in contrast to our modeling assumptions.

Finally, we compared the specific and volumetric yield achieved by the experiments in this study (at *t* = 31 h) to published values. Table 2 shows that in terms of volumetric and specific yield the pDNA production strategy for our control values is already one of the best we could find, and with 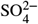 limitation it performs better than all other published methods of our knowledge. Due to the extraordinary pDNA size in this study, the plasmid copy number is just above average. However, compared to the control, the plasmid copy number of the sulfate limited process increases by 29 %.

**Table 2:**
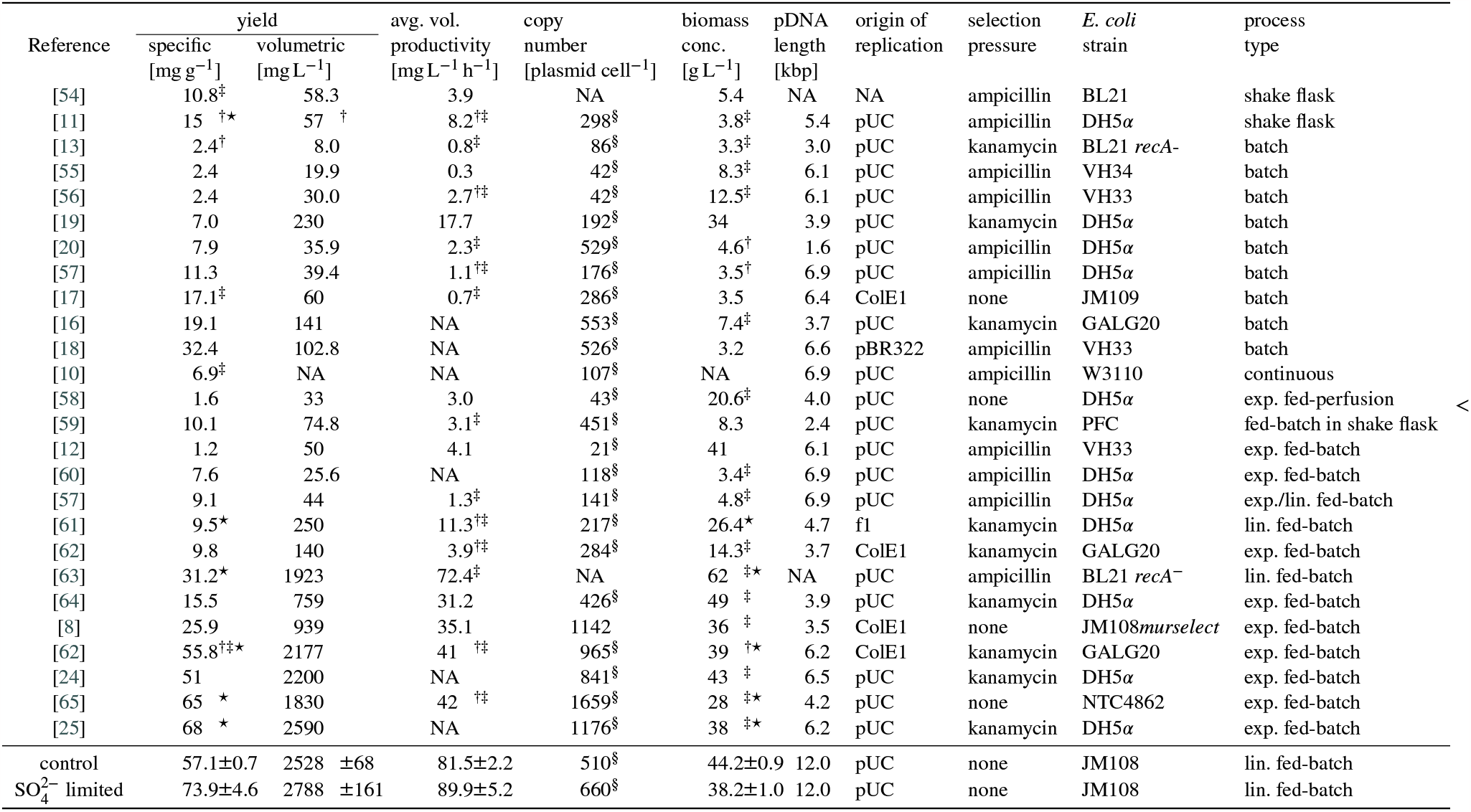
Comparison of pDNA specific yields and volumetric yields in published studies. The last two rows show the results of the experiments of this study (all pDNA conformations). Values marked with † are estimated from published figures. Values marked with ‡ are calculated from reported values. Values marked with ⋆ are converted from OD_600_ to cell dry mass (0.33 g/L/OD_600_, BNID109838 [53]). Values marked with § are estimated with 50 % GC content and an *E. coli* cell mass of 110 fg (BNID100498 [53]). Values reported as NA were not accessible.

## 4. Discussion

We aimed to improve pDNA productivity by designing a three-stage fed-batch process that separates cellular growth and production. Growth-decoupled processes are common design choices to enhance volumetric productivity in biochemical and biopharmaceutical production processes [43–46]. Especially with the advent of dynamic control in metabolic engineering that allows switching back and forth between metabolic growth and production phenotypes, interest in such (multi-stage) process designs has strongly grown [47]. While algorithms like MoVE [48] exist to identify intracellular metabolic switches, our focus here was on easily implementable medium modifications to induce these switches.

In this study, we identified twelve possible decoupling components (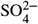, K^+^, Ca^2+^, Cl^−^, and compounds of trace elements) for the production of pDNA (Figure 2 and 3). All of them enable and regulate key functions in life [49]. Although trace elements act primarily as catalysts in enzyme systems, some of them, like copper and iron, play vital roles in energy metabolism [50]. However, we specifically selected 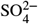 for further investigations because: (i) Sulfate is one of the six most prevalent elements in living organisms [41], which makes it comparably easy to measure and consequently determine the onset of starvation; (ii) Sulfate, in contrast to the other decoupling compounds, has a dedicated metabolic function that is well captured in the used genome-scale metabolic reconstruction *i*ML1515 of *E. coli*. [29]; (iii) Sulfate itself neither has a catalytic function nor a role in energy metabolism [51].

We simulated a three-stage fed-batch process where the transition from growth to production is triggered by the onset of sulfate starvation. Our computational model is based on two assumptions: (i) In each phase the specific pDNA production rate is constant; (ii) pDNA productivity increases upon starvation, i.e. 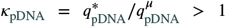. The latter aligns with experimental results of Masuda et al. [43], who reported a value of *κ*_mevalonate_ = 1.16 for mevalonate production during sulfate starvation.

In our experiments, we observed a decrease in the specific pDNA production for both the control and 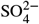 limited process (Figure 7B), challenging our assumption of a constant specific pDNA production rate. Investigating why *q*_pDNA_ decreases throughout the process will be the scope of further work. However, we implemented a time-dependent *q*_pDNA_ in additional simulations which demonstrate that the assumption of constant *q*_pDNA_ is not necessary for process improvements by 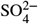 limitation (dotted lines in Figure 7 and S6) Even the optimal switching time changes by less than 1 h (Figure S7). Sulfate starvation always increases pDNA production, provided that *κ*_pDNA_ > 1, regardless of the process (batch, exponential or linear fed-batch). In fact, our data consistently shows higher specific pDNA production rates during starvation compared to control (Figure 7C). This trend reinforces and validates our core assumption of *κ*_pDNA_ > 1 where the length of starvation required to maximize volumetric productivity strongly depends on its exact value.

Maintaining high *κ*_pDNA_ during sulfate starvation is a key requirement of our design. Our predictions are based on continuously elevated levels of pDNA productivity throughout starvation. For long starvation phases, this assumption may not be feasible [52]. However, Figure 8 illustrates that this assumption is not particularly crucial. In the worst case (at *κ*_pDNA_ = 160 %), pDNA production needs to be maintained for 8.3 h to perform at least as well as a non-starved process. Our experimental data (Figure 5 and 7) demonstrate that this is indeed feasible. Interestingly, if *κ*_pDNA_ is raised beyond 160 %, the optimal starvation time increases too, but the minimally required length of pDNA production during starvation drops. This hints at a possible trade-off that may be explored in further process optimization steps.

**Figure 8:**
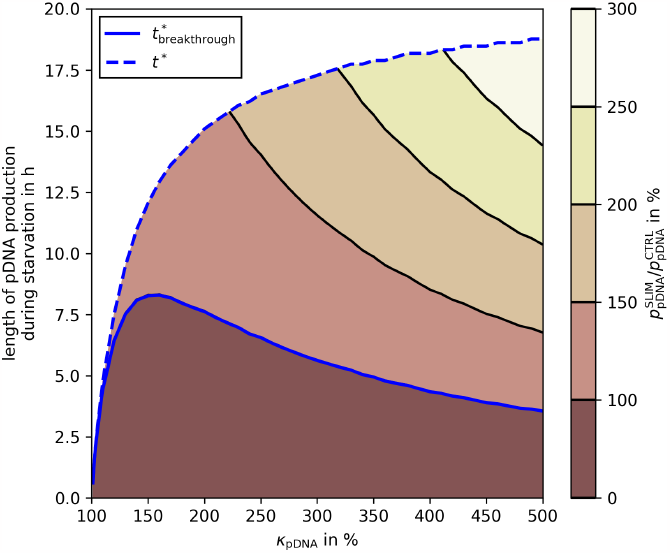
Breakthrough production length during starvation in linear fed-batch processes. If the pDNA production during starvation can be held longer than the breakthrough production length (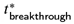 blue full line), the 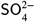 limited process outperforms a control (i.e., not starved) process. A visual explanation of 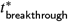 is given in Figure S4. The start of the starvation is defined by the optimal 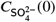 calculated in Figure 4A (red dotted line). The maximum pDNA production length during starvation is equal to the total starvation length *t*^*^ (blue dashed line). The contour colors indicate the productivity of a 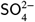 limited process compared to the control in % (color bar on the right).

In a growth-decoupled process, it is essential to, first, reach high biomass which can subsequently catalyze product formation. With our process settings (i.e., fixed final process volume), this is best achieved with a linear feeding regime (Figure S2), which quickly builds up biomass during the first few hours (compare Figure S2B and E).

We experimentally verified that a linear feeding strategy (which results in a continuously decreasing growth rate) outperforms the exponential feed even without sulfate limitation (data not shown). This is in agreement with literature which shows that a lower growth rate is preferential for pDNA production [10–12]. Interestingly a literature survey (Table 2) reveals that exponential feeding strategies are more frequently used which may explain why even our linear control process is able to outperform the majority of previously reported values. However, an ultimate comparison cannot be made as these studies used different plasmids which may significantly influence the evaluation metrics of the production process [21].

A key challenge in any growth-decoupled process is to maintain metabolic activity in non-growing metabolic states. Often a strong decrease in nutrient uptake is observed [43]. However, during sulfate starvation, glucose concentration in the reactor remained below the detection limit, indicating that cells consistently maintained glucose uptake equal to the glucose feed rate. This supports the validity of our assumption stated in Equation 5. We speculate that this may be related to the fact that (i) due to the linear feed, the specific glucose uptake already dropped to 4 % of its initial value at the onset of starvation – 81 % lower than the (already) reduced specific glucose uptake rate during sulfate starvation reported by Masuda et al. [43]; (ii) sulfate starvation retains high ATP-levels compared to other nutrient limitations [45, 67].

Consistent with maintained metabolic activity, we detected acetate accumulation during starvation (Figure S5), which is a common sign of overflow metabolism in *E. coli* [68]. However, our theoretical predictions for the maximum acetate concentration exceeded the measured values, suggesting the existence of other byproducts. Identifying these will be the focus of future research.

In the future, additional refinement of the process model could be achieved by including a term for the metabolic burden of resistance protein synthesis. An appropriate computational framework has recently been published [69]. Several studies have shown that this may significantly influence the metabolism of a producing organism [7, 8].

In both control and 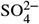 limited experiments, specific pDNA yields and concentrations dropped at the end of the bioprocess. This might be due to other limitations (e.g., the O_2_ transfer rate [70]). Therefore, we suggest stopping the process at 31 h. At that point product concentration, average volumetric productivity, and specific yield are statistically significantly up by 10 %, 10 %, and 29 %, respectively (compared to control). Considering the fraction of supercoiled pDNA, the sulfate limited process gains another 3 % points to concentration and productivity, and 4 % points to specific yield. A mechanistic interpretation of this interesting observation, however, is outside the scope of the current methodology and will be the focus of further work.

Despite using a defined, minimal medium and a comparatively large plasmid (12 kbp) our process outperforms previous reports [25] achieving an 8 % increase in volumetric yield and a 9 % boost in specific yield (Table 2). The latter is not only advantageous for higher pDNA quantities in sulfate limited experiments but, importantly, also aids in downstream processing [71, 72].

To showcase the broad applicability of sulfate limitation in biotechnological production, we explored the effect of sulfate limitation on 307 naturally secreted products of *E. coli* that are listed in the *i*ML1515 model [29]. Through lexicographic FBA with and without sulfate in the medium, we identified 83 compounds – more than 25 % of those investigated – that would benefit from sulfate limitation in their production (Figure 9). This underscores the significant potential of sulfate limited process design for a wide range of biotechnological products.

**Figure 9:**
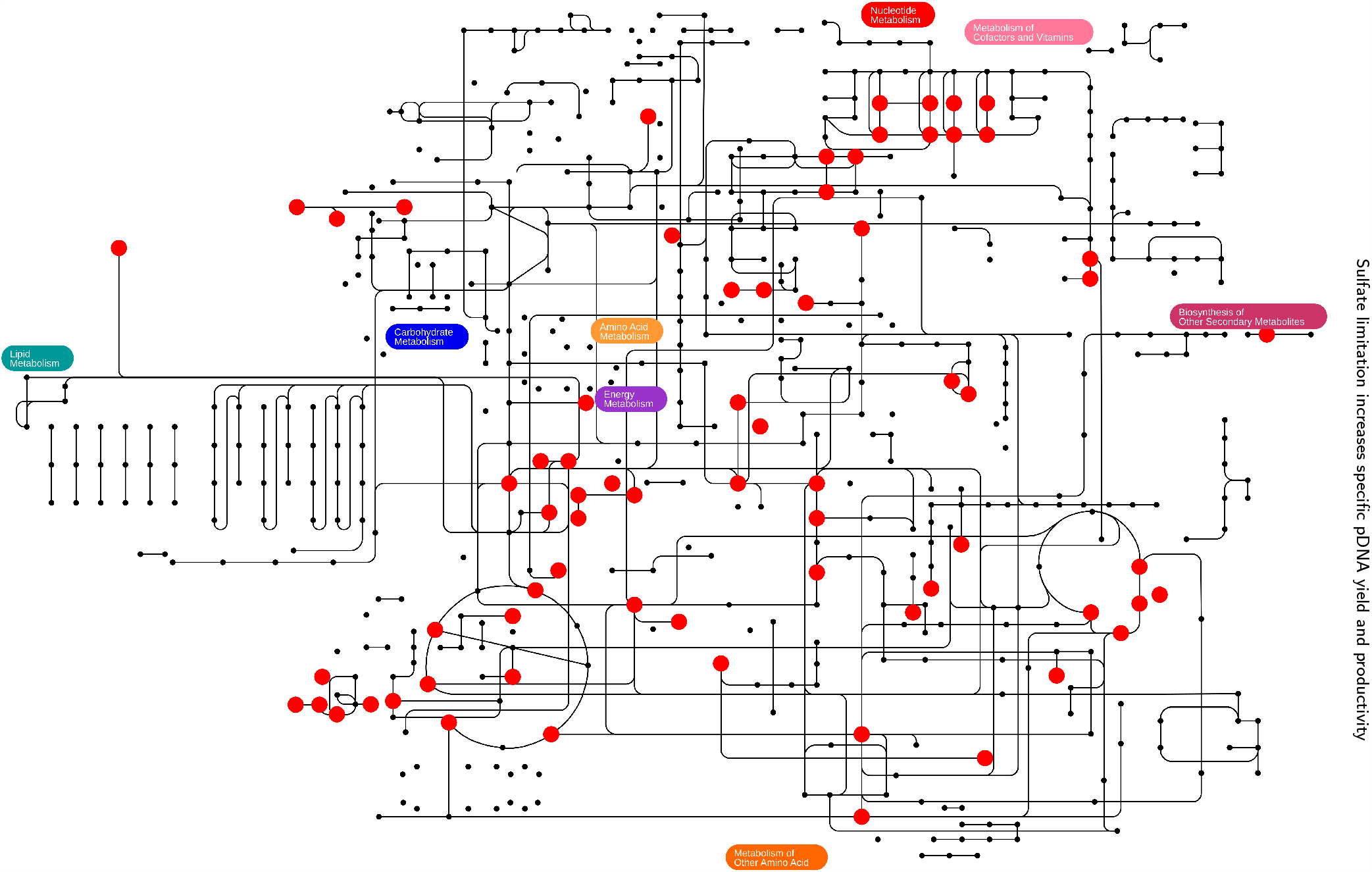
Applicability of sulfate limitation to other biotechnologcial products. Dots and lines indicate metabolites and reactions, respectively, annotated in *i*ML1515. Metabolites shown as red circles are targets for productivity improvement by sulfate limitation. The figure was created with iPath 3.0 [66]. An interactive version of the map is available at https://pathways.embl.de/selection/DfMG1nmoIuPIvOcdQSt. A full list of identified targets is shown in Table S4.

## 5. Conclusion

Based on genome-scale metabolic modeling, we have designed and successfully validated a three-stage, growth-decoupled fed-batch process for pDNA production in *E. coli*, cultivated in a minimal medium. We achieved the transition between the growth and production phases through sulfate starvation. This optimization led to statistically significant increases in key metrics: average supercoiled volumetric productivity (+13 %), specific pDNA yield (+29 %), and supercoiled specific pDNA yield (+33 %). Overall, our process achieved a specific pDNA yield of 74 mg g^−1^ and a volumetric yield of 8 g L^−1^, marking an increase of more than 8 % compared to prior reports. Importantly, our process design may be benefitial to a wide range of bio-based products of industrial significance.

## Ethics Approval and Consent to Participate

Not applicable.

## Consent for Publication

Not applicable.

## Availablility of Data and Materials

All scripts and data are available at https://github.com/Gotsmy/slim.

## Acknowledgements

Not applicable.

## Authors’ Information

^a^ Department of Analytical Chemistry, University of Vienna, Vienna, 1090, Austria. ^b^ Doctorate School of Chemistry, University of Vienna, Vienna, 1090, Austria. ^c^ enGenes Biotech GmbH, Vienna, 1190, Austria. ^d^ Baxalta Innovations GmbH, A Part of Takeda Companies, Orth an der Donau, 2304, Austria.

## CRediT authorship contribution statement

**Mathias Gotsmy:** Conceptualization, Methodology, Software, Formal Analysis, Investigation, Visualization, Writing – original draft, review and editing. **Florian Strobl:** Methodology, Formal Analysis, Investigation, Writing – review and editing. **Florian Weiß:** Methodology, Formal Analysis, Investigation, Writing – review and editing. **Petra Gruber:** Funding, Writing – review and editing. **Barbara Kraus:** Funding, Writing – review and editing. **Juergen Mairhofer:** Funding, Writing – review and editing. **Jürgen Zanghellini:** Conceptualization, Funding, Writing – original draft, review and editing.

