## Supplementary Information for "Sulfate limitation increases specific plasmid DNA yield and productivity in *E. coli* fed-batch processes"

### A. Supplementary Information

#### A.1. Supplementary Tables

| Parameter | Batch | Fed-Batch Exponential | Fed-Batch Linear | Unit |
|---|---|---|---|---|
| $C_{biomass}(0)$ | 0.02 | 3.04 | 3.04 | $g\,L^{-1}$ |
| $C_{glucose}(0)$ | 20 | 0 | 0 | $g\,L^{-1}$ |
| $C_{SO_4^{2-}}(0)$ | 0 ... 0.29 | 0 ... 3.8 | 0 ... 2.9 | $g\,L^{-1}$ |
| $C_{pDNA}(0)$ | 0 | 0 | 0 | $g\,L^{-1}$ |
| $C_{glucose}^{feed}$ | – | 330 | 330 | $g\,L^{-1}$ |
| $V(0)$ | 0.5 | 0.5 | 0.5 | L |
| $V(t_{end})$ | 0.5 | 1 | 1 | L |
| $t_{end}$ | var | 35.2 | 36.0 | h |
| $k_M$ | 2.7 [73] | – | – | $mg\,L^{-1}$ |
| $\mu$ | var | 0.1 | var | $h^{-1}$ |
| $q_{glucose}^{max}$ | 1.9 [74] | – | – | $g\,g^{-1}\,h^{-1}$ |
| $q_{pDNA}^{\mu}$ | 4.9 | 4.9 | 4.9 | $mg\,g^{-1}\,L^{-1}$ |
| $q_{ATPM}$ | 6.86 | 6.86 | 6.86 | $mmol\,g^{-1}\,h^{-1}$ |
| $r_i^{feed}$ | – | Eq. (4b) | 13.9 | $mL\,h^{-1}$ |
| $Y_{feed/biomass}$ | – | 10 | – | $mL\,g^{-1}$ |

Table S1: **Parameters for dFBA simulations.** Cells marked with "–" are not applicable and cells marked with "var" are not constant in the respective process.

| Component | Batch | Fed-Batch $SO_4^{2-}$ limited | Fed-Batch control | Unit |
|---|---|---|---|---|
| glucose $\cdot$ $H_2O$ | 10.0 | 339.9 | | g/L |
| $Na_2HPO_4 \cdot 2\ H_2O$ | 7.38 | 7.38 | | g/L |
| $KH_2PO_4$ | 5.44 | 5.44 | | g/L |
| citric acid | 2.0 | 2.0 | | g/L |
| $NH_4Cl$ | 2.0 | 0 | | g/L |
| $MgSO_4 \cdot 7\ H_2O$ | 0.24 | 3.8$^{\dagger}$ | 7.2 | g/L |
| L-proline | 0.20 | 6.0 | | g/L |
| L-isoleucie | 0.20 | 6.0 | | g/L |
| bacto yeast extract | 0.1 | 0 | | g/L |
| $MgCl_4 \cdot 6\ H_2O$ | 0 | 0.42$^{\ddagger}$ | 0 | g/L |
| trace element solution | 16.68 | 50 | | mL/L |
| thiamine-HCl (1%) | 0.10 | 3.0 | | mL/L |

Table S2: Growth media components of the experiments. Growth media were sterilized by filtration. The trace element solution was prepared in 5M HCl and contained (g/L): 4.41 $CaCl_2 \cdot 2H_2O$, 3.34 $FeSO_4 \cdot 7H_2O$, 1.43 $CoCl_2 \cdot 6H_2O$, 1.03 $MnSO_4 \cdot H_2O$, 0.15 $CuSO_4 \cdot$ 5H2O, 0.17 $ZnSO_4 \cdot 7H_2O$. † In total, 9.6 mL of a 200 g/L $MgSO_4 \cdot 7\ H_2O$ stock solution were pulsed into the bioreactor (3.2 mL at feed start, 3.2 mL at 7 h after feed start and 3.2 mL at 15 h after feed start). ‡ According to our stoichiometric model [29], the amount of $MgSO_4 \cdot 7\ H_2O$ in the batch medium alone can support more than 50 g of biomass. However, to ensure that $Mg^{2+}$ is present in excess throughout the sulfate limited process we further added $Mg^{2+}$ in the form of $MgCl_2 \cdot 6\ H_2O$.

| Process Variable | Unit | CTRL | SLIM | Change | $p$-Value |
|---|---|---|---|---|---|
| $C_{biomass}$ | g/L | 44.233 | 37.750 | −15% | 1.0000 |
| ccc fraction | | 0.878 | 0.904 | + 3% | 0.0019** |
| $M_{ccc\ pDNA}$ | g | 2.060 | 2.316 | +12% | 0.0061** |
| $C_{ccc\ pDNA}$ | g/L | 2.220 | 2.519 | +13% | 0.0052** |
| $p_{ccc\ pDNA}$ | g/L/h | 0.072 | 0.081 | +13% | 0.0052** |
| $Y_{ccc\ pDNA/biomass}$ | g/g | 0.050 | 0.067 | +33% | 0.0001*** |
| $M_{pDNA}$ | g | 2.346 | 2.564 | + 9% | 0.0291* |
| $C_{pDNA}$ | g/L | 2.528 | 2.788 | +10% | 0.0243* |
| $p_{pDNA}$ | g/L/h | 0.082 | 0.090 | +10% | 0.0243* |
| $Y_{pDNA/biomass}$ | g/g | 0.057 | 0.074 | +29% | 0.0005*** |

Table S3: Average values of several process variables of interest and their $p$-values compared to the control at 31 h (one-sided $t$-test). ccc pDNA corresponds to the supercoiled fraction of plasmid DNA measured.

| Compound | max. $q^\mu$ | max. $q^*$ | Compound | max. $q^\mu$ | max. $q^*$ |
|---|---|---|---|---|---|
| mththf† | 0.000 | 9.610 | (R)-Glycerate | 0.000 | 21.367 |
| (R)-Propane-1,2-diol | 0.000 | 13.978 | (S)-Propane-1,2-diol | 0.000 | 13.839 |
| ca4colipa† | 0.000 | 0.183 | 1,5-Diaminopentane | 0.000 | 7.723 |
| 2-Oxoglutarate | 0.000 | 12.762 | 3-Hydroxypropanoate | 0.000 | 11.744 |
| 4-Aminobutanoate | 0.000 | 11.913 | 4-aminobenzoyl-glutamate | 0.000 | 4.533 |
| 5-Dehydro-D-gluconate | 0.000 | 10.529 | Acetaldehyde | 0.000 | 23.501 |
| Acetate | 0.000 | 28.795 | Adenine | 0.000 | 10.334 |
| Adenosine | 0.000 | 5.330 | Agmatine | 0.000 | 8.340 |
| Allantoin | 0.000 | 13.197 | Citrate | 0.000 | 11.530 |
| Cytidine | 0.000 | 6.131 | D-Alanine | 0.000 | 13.515 |
| D-Alanyl-D-alanine | 0.000 | 6.758 | D-Gluconate | 0.000 | 10.077 |
| D-Glyceraldehyde | 0.000 | 17.742 | D-Lactate | 0.000 | 20.000 |
| Dihydroxyacetone | 0.000 | 18.511 | Enterochelin | 0.000 | 1.704 |
| Ethanol | 0.000 | 20.000 | Ethanolamine | 0.000 | 10.830 |
| fad† | 0.000 | 1.769 | FMN C17H19N4O9P | 0.000 | 2.700 |
| Fe(III)dicitrate | 0.000 | 5.765 | Fe-enterobactin | 0.000 | 1.704 |
| Formate | 0.000 | 102.256 | Fumarate | 0.000 | 18.482 |
| Glycerol 3-phosphate | 0.000 | 9.606 | Glycerol | 0.000 | 15.637 |
| Glycerophosphoglycerol | 0.000 | 5.675 | Glycine | 0.000 | 27.012 |
| Glycolate C2H3O3 | 0.000 | 28.795 | Guanine | 0.000 | 10.708 |
| Guanosine | 0.000 | 5.395 | Hexanoate (n-C6:0) | 0.000 | 7.500 |
| Hypoxanthine | 0.000 | 10.956 | Indole | 0.000 | 5.412 |
| Inosine | 0.000 | 5.459 | L-Alanine | 0.000 | 19.860 |
| L-Arginine | 0.000 | 8.274 | L-Asparagine | 0.000 | 15.935 |
| L-Aspartate | 0.000 | 17.975 | L-Glutamate | 0.000 | 11.815 |
| L-Histidine | 0.000 | 8.063 | L-Homoserine | 0.000 | 13.002 |
| L-Idonate | 0.000 | 10.077 | L-Isoleucine | 0.000 | 7.240 |
| L-Lactate | 0.000 | 17.376 | L-Leucine | 0.000 | 7.514 |
| L-Lysine | 0.000 | 7.723 | L-Malate | 0.000 | 18.482 |
| L-Phenylalanine | 0.000 | 5.428 | L-Proline | 0.000 | 10.019 |
| L-Serine | 0.000 | 20.626 | L-Threonine | 0.000 | 12.325 |
| L-Tryptophan | 0.000 | 4.465 | L-Tyrosine | 0.000 | 5.635 |
| L-Valine | 0.000 | 9.930 | LalaDgluMdap† | 0.000 | 3.174 |
| LalaDgluMdapDala† | 0.000 | 2.739 | anhgm† | 0.000 | 2.669 |
| O-Acetyl-L-serine | 0.000 | 11.800 | Ornithine | 0.000 | 9.478 |
| nlipa† | 0.000 | 0.397 | Protoheme C34H30FeN4O4 | 0.000 | 1.301 |
| Putrescine | 0.000 | 9.565 | Pyruvate | 0.000 | 23.057 |
| g3p† | 0.000 | 6.361 | Succinate | 0.000 | 17.137 |
| Thymidine C10H14N2O5 | 0.000 | 5.053 | Thymine C5H6N2O2 | 0.000 | 10.311 |
| Uracil | 0.000 | 15.672 | Urea CH4N2O | 0.000 | 28.228 |
| Uridine | 0.000 | 6.510 | Xanthine | 0.000 | 11.921 |
| Xanthosine | 0.000 | 5.758 | | | |

Table S4: List of potential targets for process optimization with sulfate limitation. We used lexicographic FBA (Table 1) to calculate maximal theoretical production rates during sulfate starvation (max. $q^*$) and sulfate excess (max. $q^\mu$) in mmol $g^{-1}$ $h^{-1}$. Compounds marked with † are displayed with their BiGG ID [75] to shorten the name. Previously described sulfate limitation target mevalonate is not naturally synthesized in *E. coli* [43] and, thus, not present in the list.

#### A.2. Supplementary Figures

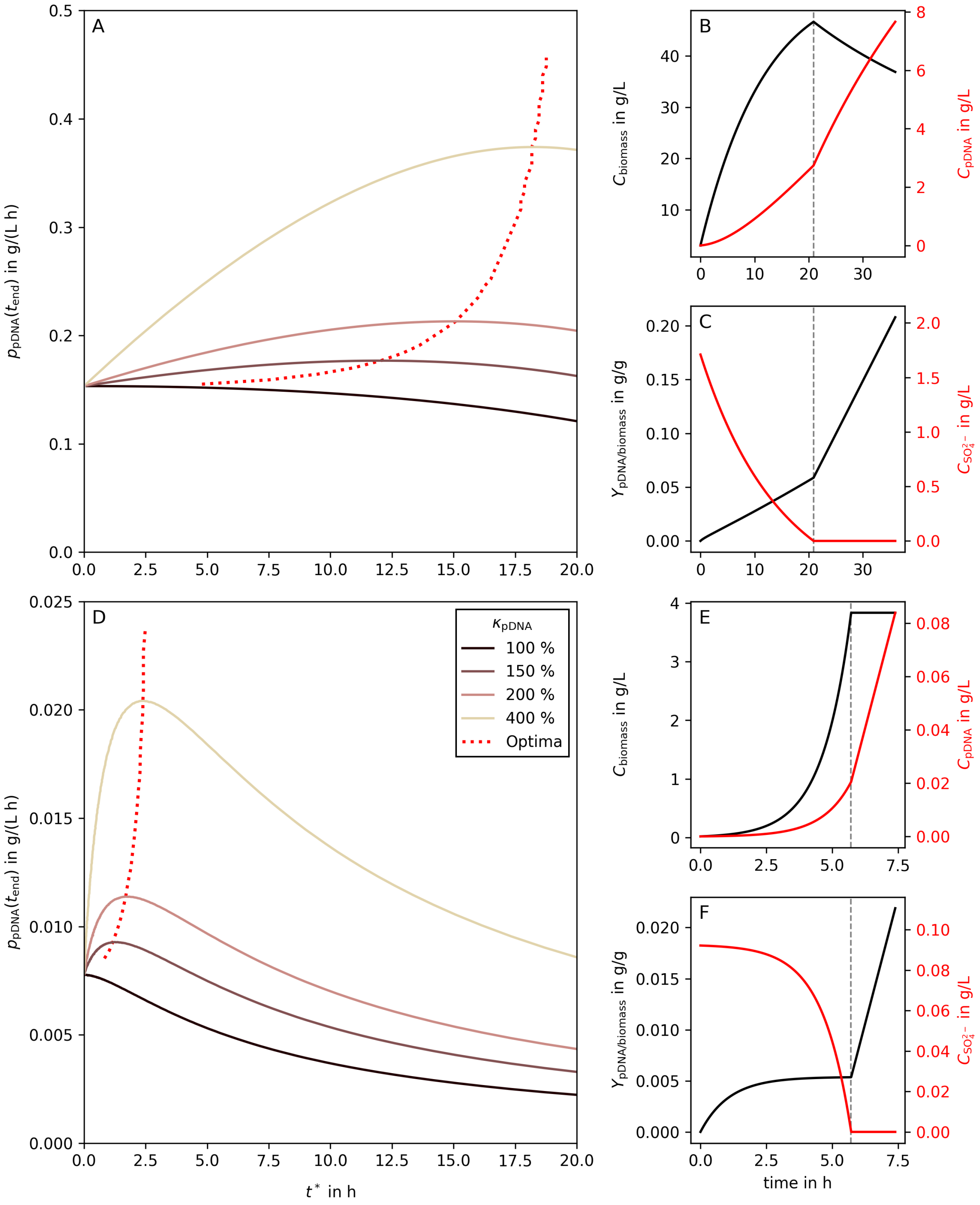


Figure S1: **Predicted optimal timing in a $SO_4^{2-}$ limited linear fed-batch (panels A, B, C) and batch (D, E, F) process.** Panels **A** and **D** show average volumetric productivities of pDNA production ($\mathrm{g\,L^{-1}\,h^{-1}}$) as a function of the length of $SO_4^{2-}$ starvation in the respective process ($t^*$). The full lines represent different levels of increased pDNA production during starvation as percentages of pDNA production rate during biomass growth ($\kappa^*_{\mathrm{pDNA}}$). The dotted line indicates the location of the optima for $\kappa^*_{\mathrm{pDNA}}$ between 0 and 500 %. Panels **B**, **C** and **E**, **F** show the process curves of metabolites of interest in the optimal respective process for $\kappa^*_{\mathrm{pDNA}} = 200\,\%$. The gray dashed lines indicate the switching time point between the growth and production phases.

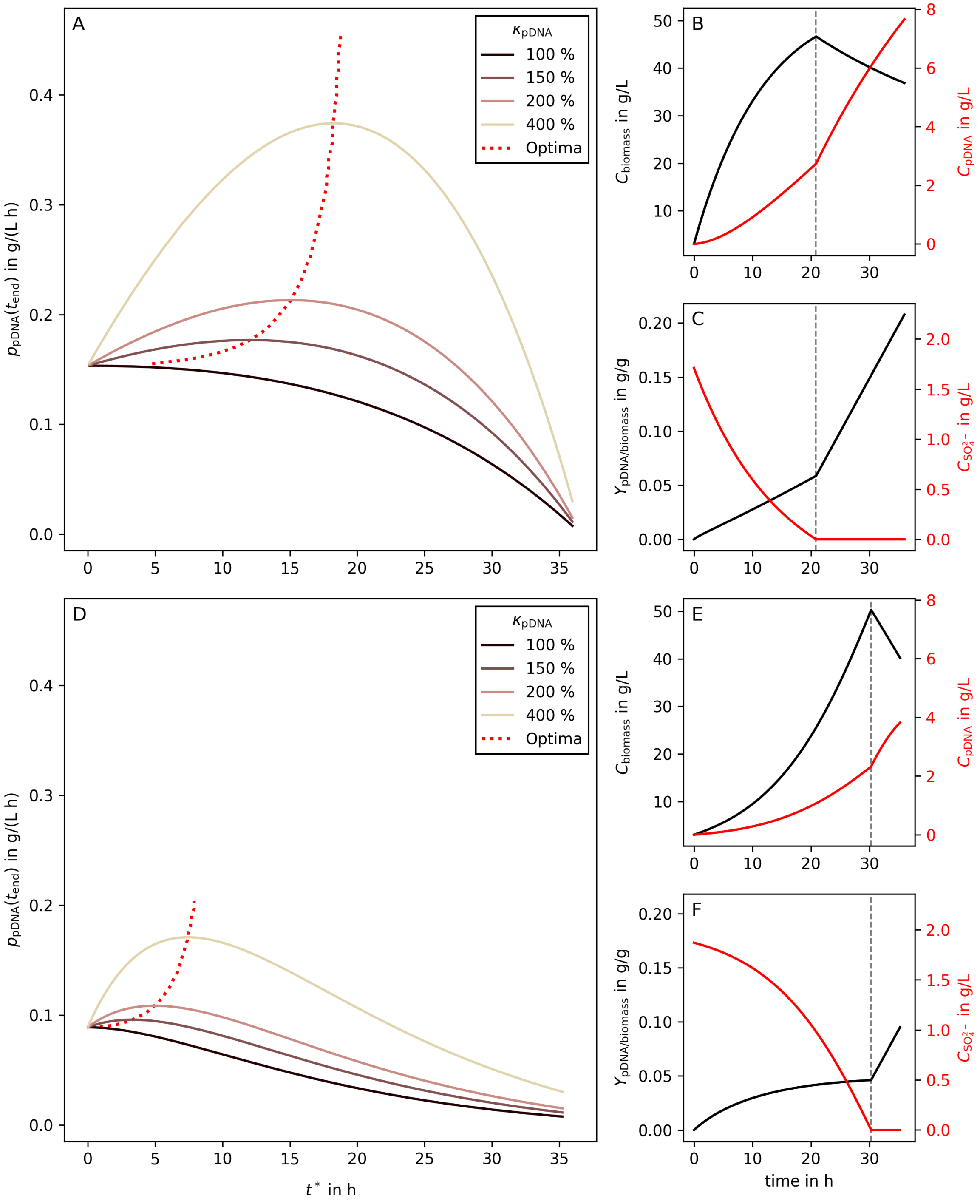


Figure S2: **Predicted optimal timing in a $SO_4^{2-}$ limited linear (panels A, B, C) and exponential (D, E, F) fed-batch process.** Panels **A** and **D** show average volumetric productivities of pDNA production ($\mathrm{g\,L^{-1}\,h^{-1}}$) as a function of the length of $SO_4^{2-}$ starvation in the respective fed-batch process ($t^*$). The full lines represent different levels of increased pDNA production during starvation as percentages of pDNA production rate during biomass growth ($\kappa^*_{\mathrm{pDNA}}$). The dotted line indicates the location of the optima for $\kappa^*_{\mathrm{pDNA}}$ between 0 and 500 %. Panels **B**, **C** and **E**, **F** show the process curves of metabolites of interest in the optimal respective process for $\kappa^*_{\mathrm{pDNA}} = 200\,\%$. The gray dashed lines indicate the switching time point between the growth and production phases.

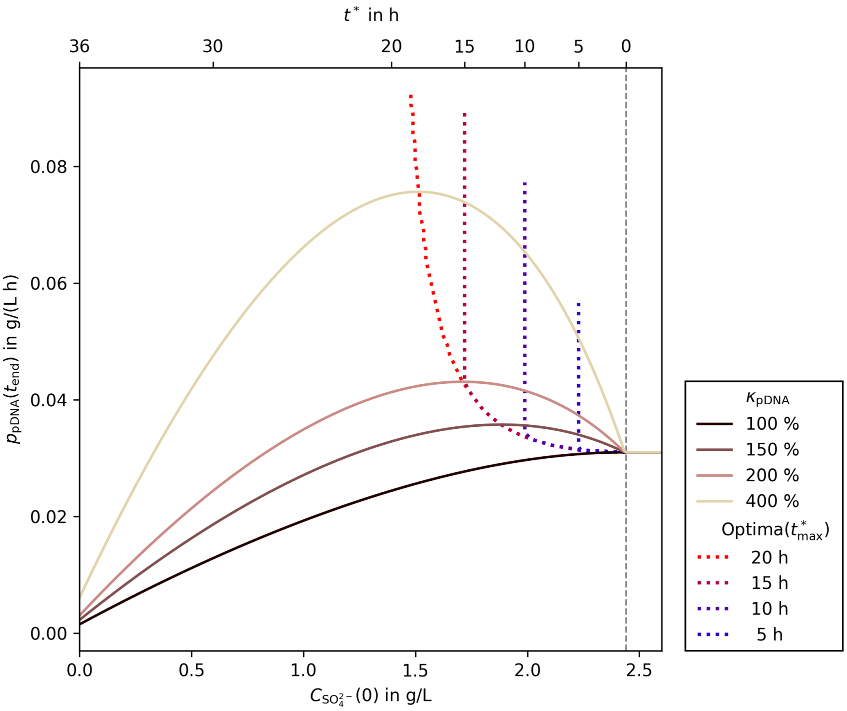


Figure S3: **Predicted optimal timing in a $SO_4^{2-}$ limited linear fed-batch process with different maximum starvation lengths.** For models right of the gray dashed line no $SO_4^{2-}$ limitation occurs. The colored dotted lines illustrate the optima of different maximum starvation phase lengths. The colored full lines illustrate different maximal pDNA production rates in the starvation phase.

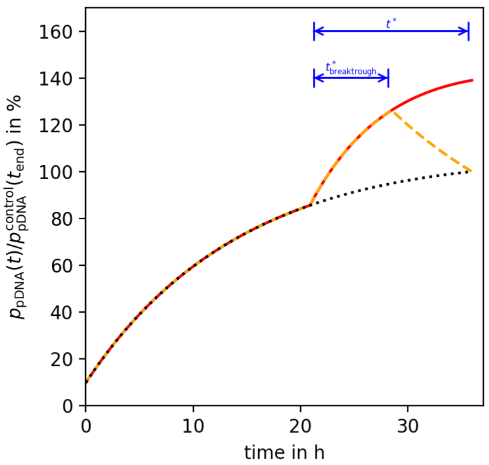


Figure S4: **Visualization explanation of $t^*_{breakthrough}$.** The black dotted line represents a non $SO_4^{2-}$ limited control process. The processes represented by the red full and the orange dashed line start their $SO_4^{2-}$ starvation at the optimal time point as calculated in Figure 4 ($\kappa_{pDNA} = 200\,\%$). The difference between them is that for the red full line, the pDNA production is held constant throughout the whole length of the starvation ($t^*$). In contrast, the orange dashed line describes $p_{pDNA}$ when pDNA production occurs only in the first $t^*_{breakthrough}$ hours of starvation and is zero thereafter. $t^*_{breakthrough}$ describes the minimal length of pDNA production during starvation that leads to equal or better final productivities ($p_{pDNA}(t_{end})$) than the control.

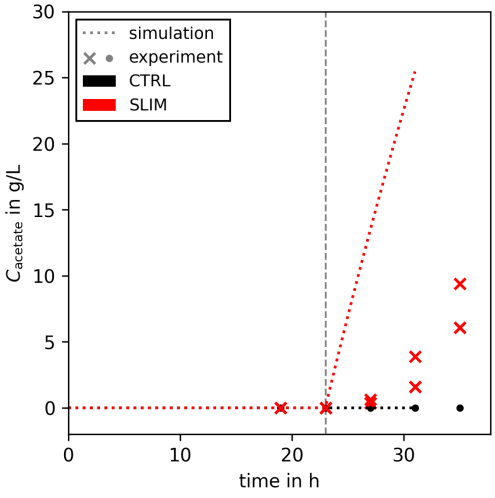


Figure S5: **Acetate accumulation during control and $SO_4^{2-}$ limited processes.** The concentration of acetate over time was measured for one and two replicate(s) of CTRL and SLIM, respectively. Glucose concentrations were measured at the same time points and stayed at 0 g $L^{-1}$ throughout the process (data not shown). Additionally, we performed dFBA simulations with time dependent $q_{pDNA}$ (dotted lines, Figure 7B) and added the maximization of acetate flux at the end of the lexicographic objectives (Table 1). Acetate concentration according to the simulations are visualized as dotted lines.

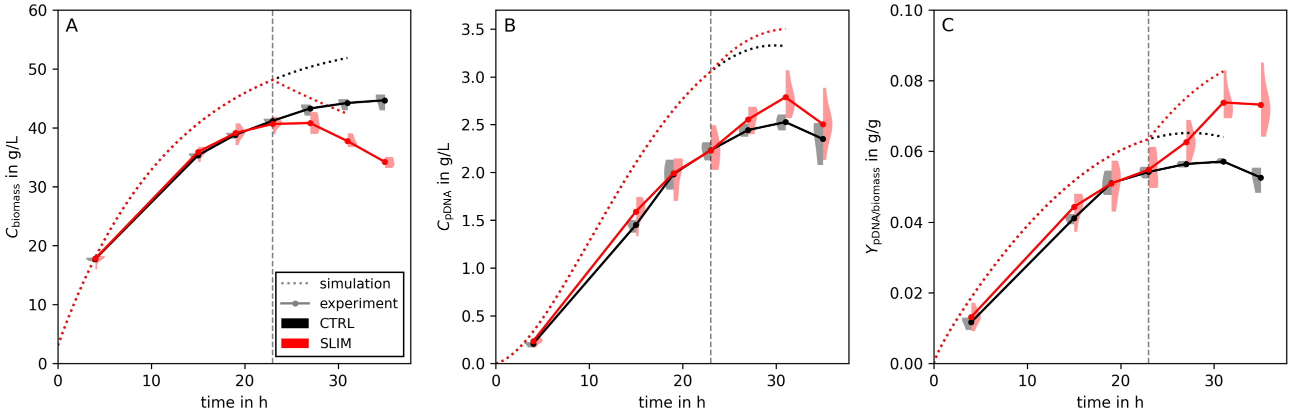


Figure S6: **Control and sulfate limited process with time-dependent $q_{pDNA}$.** A time-dependent, linearly declining $q_{pDNA}^{CTRL}(t)$ rate (black dotted line, Figure 7B) was fitted to the experimental data (black circles, Figure 7B). The rate of $q_{pDNA}^{SLIM}(t)$ was shifted in parallel to minimize the error of the flux ration at $q_{pDNA}$ starved experimental data points (i.e., at $t = 27$ and 31 h, blue dotted line, Figure 7C). Subsequently, we performed dFBA. Results for biomass and pDNA concentration, and specific yield are plotted in panels A, B, and C, respectively. The simulation highlights that the assumption of constant $q_{pDNA}$ is not necessary for process improvements by $SO_4^{2-}$ limitation.

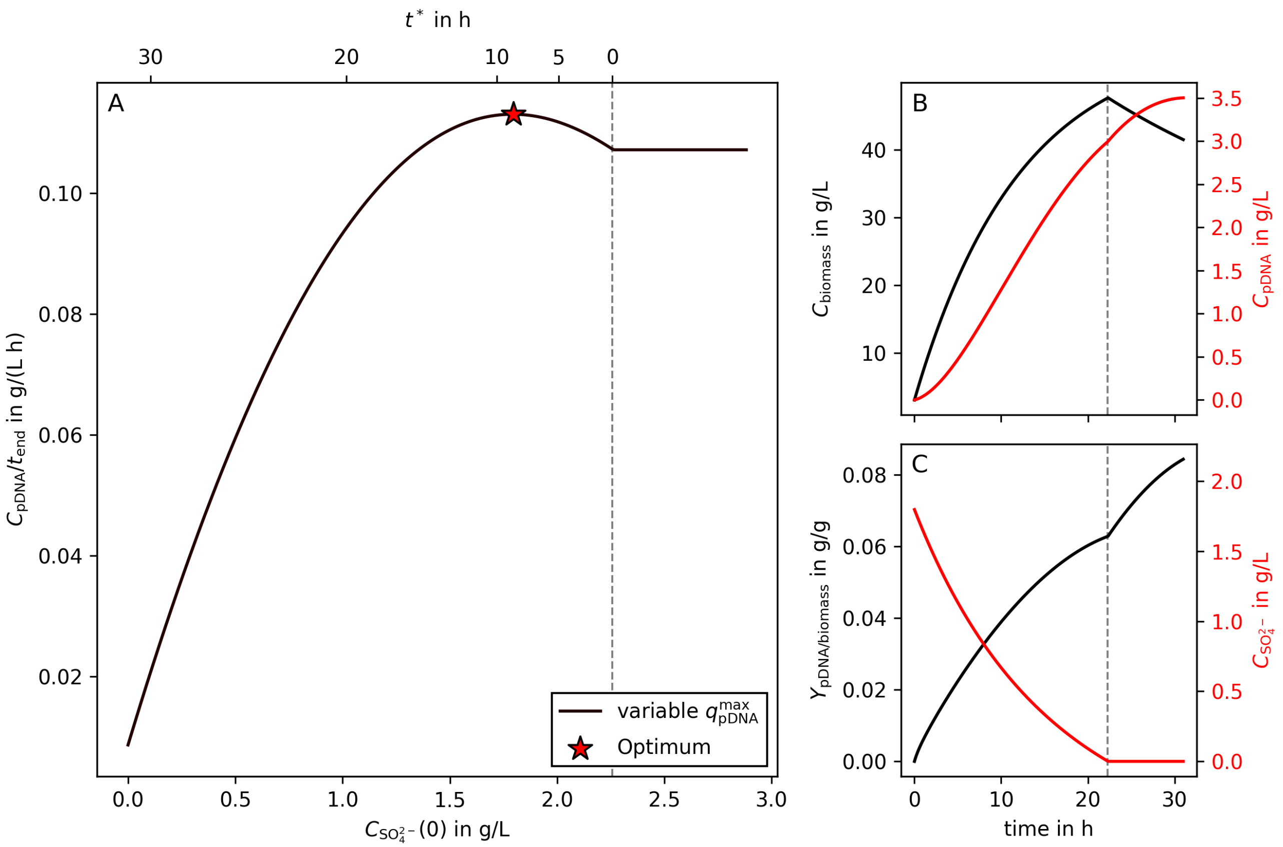


Figure S7: **Predicted optimal timing in a $SO_4^{2-}$ limited linear fed-batch process with variable $q_{pDNA}$.** $q_{pDNA}$ rates were fitted to CTRL and SLIM experimental values retroactively (Figure 7C). Panel **A** shows average volumetric productivities of pDNA production ($g\,L^{-1}\,h^{-1}$) as a function of the initial $SO_4^{2-}$ concentration. The second X-axis of panel **A** (top) illustrates the length of the sulfate starved process phase ($t^*$). For all modeled linear fed-batch processes, right of the gray dashed line, no $SO_4^{2-}$ limitation occurred. Panels **B** and **C** show the process curves of metabolites of interest in the optimal process. The gray dashed lines indicate the switching time point between the growth and production phases at $t = 22.3$ h instead of the initially predicted 23 h with constant $q_{pDNA}$ (Figure 4).

#### A.3. Supplementary Notes

##### *A.3.1. Optimal Sulfate Limited Batch Process*

*Methods* Initial conditions (Table S1) represent a realistic batch process in a small bioreactor. Glucose uptake rate, $q_{\text{glucose}}(t)$, was modeled as Michaelis-Menten kinetics,

$$q_{\text{glucose}}(t) = \frac{q^{\max}_{\text{glucose}} C_{\text{glucose}}(t)}{k_{\text{M}} + C_{\text{glucose}}(t)} \tag{S1}$$

with $C_{\text{glucose}}$ denoting the current glucose concentration in the medium, and two constants ($q^{\max}_{\text{glucose}}, k_{\text{M}}$). Simulations terminated once the glucose concentration dropped below the level of what was required for ATP maintenance. Initial sulfate concentration was varied in 301 equidistant steps to search for a volumetric pDNA productivity optimum.

*Results* We simulated sulfate limited batch processes using the initial values listed in Table S1 and analyzed the impact of the length of the pDNA production phase (induced by sulfate starvation) on the average volumetric productivity (Figure S1D).

We observe distinct maxima in the average volumetric productivity when $\kappa_{\text{pDNA}} > 1$ (red dotted line, Figure S1D), i.e. when the pDNA production rate is enhanced. With increasing $\kappa_{\text{pDNA}}$, the maximum productivity increases and moves towards smaller values of $C_{\text{SO}_4^{2-}}(0)$. The initial sulfate concentration (not shown) is proportional to the length of the growth phase. The larger $C_{\text{SO}_4^{2-}}(0)$, the longer the growth phase and the shorter the starvation (i.e., pDNA production) phase ($t^*$), and vice versa. Thus, Figure S1D mirrors the trade-off between producing a sufficiently high cell density that catalyzes the synthesis of pDNA, and actual pDNA production [42]. For large improvements, i.e., $\kappa_{\text{pDNA}} \gtrapprox 3$, the optimal length of the starvation phase is roughly 2.5 h and becomes virtually independent of $\kappa_{\text{pDNA}}$.

As an example, the optimal productivity for $\kappa_{\text{pDNA}} = 2$ (corresponding to $q^*_{\text{pDNA}} = 9.9\,\text{mg}\,\text{g}^{-1}\,\text{h}^{-1}$) is reached by adding $0.092\,\text{g}\,\text{L}^{-1}$ $\text{SO}_4^{2-}$ into the batch medium. This results in a 1.7 h starvation phase, which increases the average volumetric productivity by 47 %. The time-dependent concentrations of this process are shown in Figure S1E and F.
